# Sustaining smallholder banana production in Banana Bunchy Top Disease endemic landscapes: integrating clean seed, roguing, and farmer training

**DOI:** 10.64898/2026.04.15.718704

**Authors:** Renata Retkute, Aman Bonaventure Omondi, Martine Zandjanakou-Tachin, Ulrich Roland Agoi, Charles Staver, P. Lava Kumar, John E. Thomas, Christopher A. Gilligan

## Abstract

Banana bunchy top virus (BBTV) continues to threaten smallholder livelihoods and food security across sub-Saharan Africa. While clean-seed programmes are widely promoted, their long-term effectiveness is often compromised by rapid reinfection in endemic landscapes. We developed an integrated framework combining spatially explicit, stochastic epidemiological modelling with additional cost–benefit analysis and use of socio-behavioural data to evaluate strategies for stabilising production. Our results demonstrate that frequent monthly inspections and accurate symptom detection are essential for disease suppression. Crucially, the economic analysis reveals that prioritizing diagnostic competence as an economic asset is necessary: improving detection efficiency can more than double farmer net revenue under realistic market conditions. Socio-behavioural findings further confirm that a farmer’s ability to correctly recognise symptoms is the strongest predictor of roguing adoption, far outweighing demographic characteristics. These results provide quantifiable guidance for disease management and highlight that the sustainability of clean-seed interventions hinges on shifting policy from simple seed replacement to investing in farmer diagnostic capacity. Strengthening this local surveillance capability transforms informal seed systems into resilient, durable tools for safeguarding household nutrition and regional food security.

## 1 Introduction

Banana bunchy top disease (BBTD), caused by the banana bunchy top virus (BBTV), has been severely affecting smallholder banana and plantain fields in sub-Saharan Africa (SSA) since it was first detected in the 1950s [1]. Over the last three decades, BBTV has spread rapidly across SSA, with expanding fronts in East, West and Southern Africa [1–8]. This spread is particularly concerning because banana and plantain (Musa spp.) are among the region’s most important food crops, serving as both a staple food and a key source of income, with over 35 million tonnes grown annually in diverse production systems [9]. Moreover, banana cultivation across sub-Saharan Africa encompasses a rich diversity of locally adapted cultivars. For example, a survey across Rwanda, Burundi, and eastern Democratic Republic of Congo recorded *>* 90 distinct cultivars [10]. The region is also a major secondary centre of banana diversity, encompassing East African Highland bananas and plantains [11], both of which are threatened by BBTD spread.

Banana bunchy top virus is transmitted primarily by the banana aphid Pentalonia nigronervosa and through infected planting material [12]. Infection leads to distinctive symptoms, including a characteristic striping resembling a Morse code pattern on emerging leaves, leaf distortion, and severe stunting that produces the characteristic bunchy top appearance, culminating in distorted or absent bunches [13]. Consequently, BBTV induces yield collapse, with losses often reaching 100% [13–15].

Breeding for resistance has successfully addressed other banana diseases and pests, including Fusarium wilt (*Fusarium oxysporum* f. sp. *Cubense*) race 1 and tropical race 4, banana weevils (C*osmopolites sordidus (Germar)*), banana bacterial wilt (*Xanthomonas vasicola pv. musacearum*), black leaf streak (*Pseudocercospora fijiensis*) [16]. For BBTD, however, no cultivated varieties of banana and plantain currently demonstrate durable field resistance [17, 18]. Partial resistance – potentially useful in emerging genetic interventions, including RNAi-based resistance [19] and genome editing approaches [20, 21] – has been found in the wild ancestor *Musa balbisiana*. Nevertheless, a set of BBTD-resistant varieties that can meet the market and consumption diversity of SSA remains a remote and distant option [22].

In the absence of resistant varieties, access to clean planting material is critical. Formal commercial systems, such as micropropagation of banana, which can generate virus-free planting material, are often limited, unaffordable, or logistically inaccessible [23]. Therefore, most farmers rely on on-farm vegetative propagation using suckers from underground rhizomes [24–26]. Although this approach is a traditional practice and a key contributor to the sustainability and gradual expansion of banana plantations at minimal cost, it also perpetuates the risk of BBTV infection and the spread of infected planting materials both in BBTV-endemic landscapes and to unaffected areas – especially when farmers inadvertently select and move infected, often pre-symptomatic, suckers between fields. While yield collapse is an immediate impact, the local spread of BBTV from field to field progressively diminishes the availability of virus-free or low-risk suckers for planting new fields, while simultaneously increasing the risk of selecting symptomless infected suckers. This vulnerability is exacerbated by the prevailing seed system: the farmer-to-farmer sucker supply system is extremely low-cost and well adapted to maintaining the wide cultivar diversity characteristic of smallholder production systems, particularly in East, Central and West Africa; however, it is highly vulnerable to sucker-borne diseases such as BBTD [25].

Insights into the specific epidemiological variables driving BBTD spread, yield loss, and sucker infection have been applied in different production systems to pilot the recovery of banana in BBTD-endemic landscapes [27]. Subsequent studies have examined varietal susceptibility [17, 28] and clean-seed deployment [29, 30], among other topics. Together, these research and field advances provide the basis for epidemiological modelling combined with cost-benefit and socio-behavioural analyses to assess how integrated strategies could sustain production under endemic conditions. In theory, frequent inspection and rapid removal of symptomatic plants can suppress BBTV spread [31–33], but in practice the effectiveness of such strategies depends critically on implementation constraints, including the timeliness, consistency, and reliability of disease detection. This is especially relevant in informal seed systems, where farmer-to-farmer exchange remains the dominant pathway for sourcing planting material, and reliance on an external seed supply is rarely feasible. Without robust on-farm monitoring, clean-seed programmes risk only short-term success because of rapid reinfection [27].

To explore these challenges, we developed an integrated framework that bridges the gap between theoretical disease suppression and the practical realities of smallholder implementation. By combining stochastic, spatially explicit epidemiological modelling with economic cost–benefit analysis and socio-behavioural survey data, we quantify the specific conditions under which clean-seed introduction, on-farm sucker propagation, and systematic roguing can successfully stabilise production in endemic landscapes. This multifaceted approach allows us to address three critical knowledge gaps: (i) determining the surveillance and management intensity required to suppress virus prevalence and sustain the production of disease-free planting material; (ii) identifying the economic conditions and thresholds under which intensive surveillance and roguing become robustly profitable for smallholders; and (iii) uncovering the socio-behavioural factors, such as diagnostic competence, that directly influence the adoption of these practices in the field. Ultimately, by integrating these epidemiological, economic, and behavioural insights, we provide a comprehensive, quantitative framework for transforming vulnerable seed systems into durable management tools that safeguard banana production as a cornerstone of daily nutrition and regional food security.

## 2 Results

### 2.1 Epidemiological outcomes under varying inspection regimes

We simulated the spread of BBTV over a four-year horizon across a range of inspection frequencies and detection probabilities. We explored detection probabilities from 10% to 100% under four inspection regimes: annual, biannual, quarterly, and monthly. Model outputs were evaluated using three metrics: BBTD prevalence; the proportion of planted BBTV-free suckers (%); and total number of banana bunches produced by healthy plants (Figure 1). For each parameter combination, we performed 1,000 stochastic simulation runs.

**Fig. 1.**
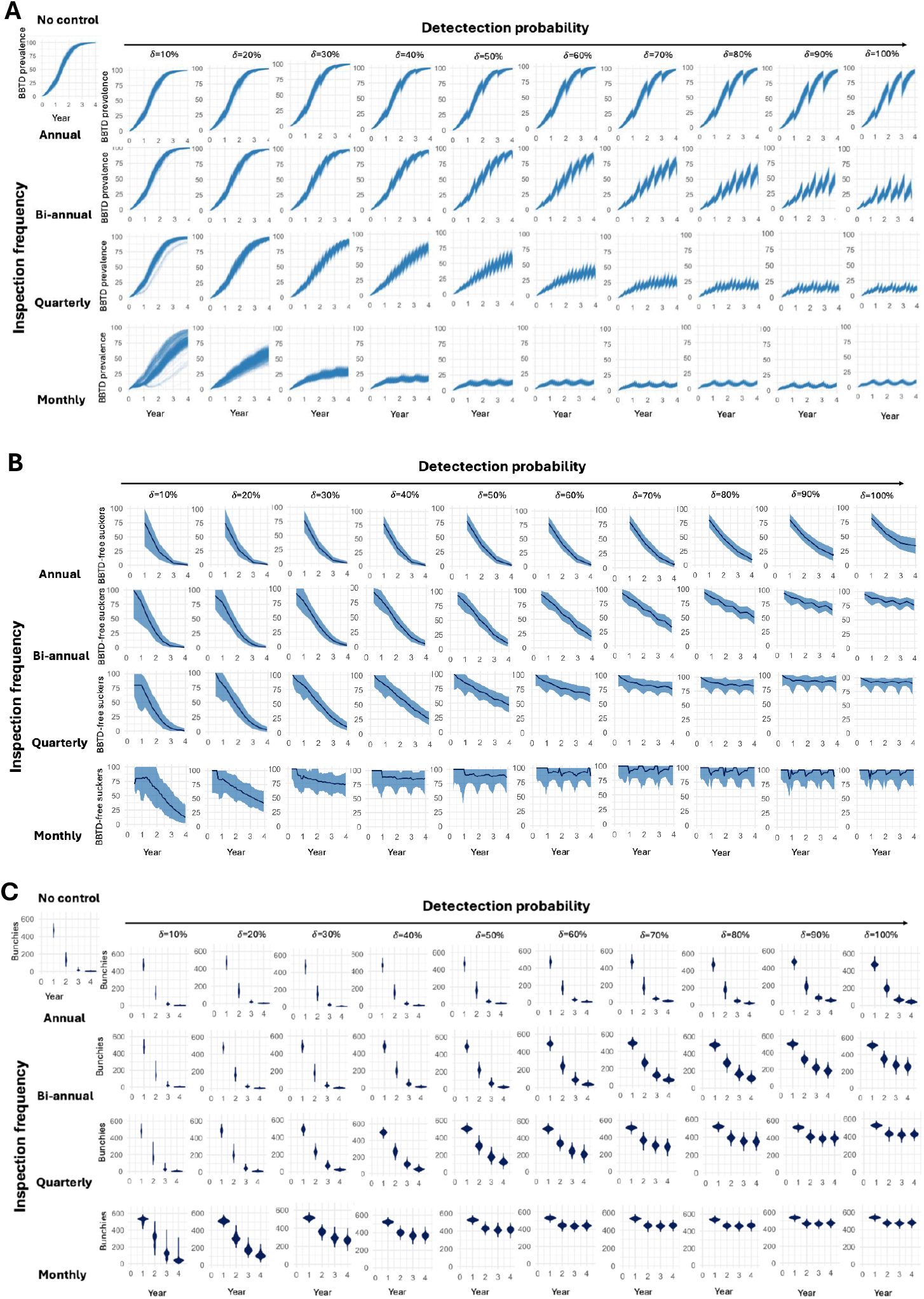
Impact of inspection frequency and detection probability over four years on: (A) BBTD prevalence; (B) % of planted BBTV-free suckers; and (C) number of banana bunches. Each panel shows results from 1,000 individual simulations. The top-left panel in A and C illustrate uncontrolled spread. Rows correspond to inspection intervals: annual (12 months), biannual (6 months), quarterly (3 months), and monthly (1 month), while columns correspond to detection probabilities (10–100%).

Model simulations demonstrated that BBTD prevalence was strongly influenced by both inspection frequency and detection probability (Figure 1A). In the absence of control (top-left panel), all simulations converged to *∼*100% BBTD prevalence by year four. More frequent inspections and higher detection probabilities consistently reduced mean, minimum, and maximum prevalence, whereas infrequent inspections allowed extensive disease spread. Annual surveillance resulted in very high prevalence across all detection probabilities. Median prevalence at four years ranged from 66% (at 100% detection) to nearly 100% (at 10–30% detection). Even with perfect detection, prevalence remained substantial (median 66%, minimum 49%). Under semi-annual inspections, prevalence was also high but showed greater sensitivity to detection probability: median prevalence ranged from 91% (50% detection) to 99% (10% detection), and only with detection *≥*60% did median prevalence fall below 90%. Quarterly inspections further reduced prevalence, but still required high detection efficiency; with detection *≥*80%, median prevalence dropped below 10%, whereas at 30% detection it remained around 60%. Monthly surveillance produced the lowest prevalence across all scenarios. At detection probabilities *≥*60%, median prevalence fell below 5%, and even at 30% detection median prevalence was approximately 31%. These results indicate that monthly inspections can effectively suppress BBTV even with moderate detection probability, whereas less frequent inspections require very high detection efficiency to achieve substantial control.

Simulation outputs showed that the proportion of BBTV-free suckers planted over time depended strongly on both inspection interval and detection probability as well (Figure 1B). Infrequent surveillance led to a rapid decline in clean planting material, while shorter inspection intervals helped preserve a higher proportion of disease-free suckers. With annual inspections, the fraction of BBTV-free suckers dropped to very low levels by year four across all detection scenarios, indicating limited capacity to maintain clean seed systems. Semi-annual inspections offered only modest improvement, with substantial losses of healthy planting material unless detection probabilities were high. Under quarterly inspections, outcomes improved markedly, particularly at higher detection levels, although low detection probabilities still resulted in pronounced declines. Monthly inspections consistently maintained the highest proportion of BBTV-free suckers, even when detection was imperfect, highlighting their effectiveness in slowing the degeneration of clean planting material.

Model simulations further showed that total banana bunch production closely tracked patterns in disease prevalence and the availability of healthy planting material (Figure 1C). Systems with infrequent inspections experienced substantial declines in bunch production, reflecting widespread infection and loss of productive plants. Under annual and semi-annual surveillance, bunch numbers remained low across most detection probabilities, with only limited gains achieved at high detection efficiency. Quarterly inspections improved productivity, particularly when detection probabilities were high, but yields were still constrained under moderate to low detection. In contrast, monthly inspections consistently resulted in the highest bunch production across all scenarios, sustaining strong yields even when detection was imperfect.

### 2.2 Quantitative characterisation of the production landscape

To establish the baseline context for BBTV management, we analysed national and field-level data on banana and plantain production, as well as producer prices. Clear long-term trends emerged in estimated bunch weight for banana and plantain over the period from 1961 to 2024 across sub-Saharan Africa (Fig. 2 A-B). Banana bunch weight showed a strong upward trajectory, increasing more than twofold from approximately 7 kg in the early 1960s to over 16 kg by 2023. Despite this overall improvement, substantial variation persisted among countries, with some maintaining low and stable bunch weights while others experienced marked increases or consistently high values. Plantain bunch weight followed a more moderate and variable trajectory. Regional averages rose from roughly 4.5–5 kg in the 1960s to a peak of approximately 7.5–8 kg in the mid-2010s, followed by a slight decline to just above 7 kg by 2024. Compared with banana, plantain exhibited weaker long-term gains and greater interannual variability, particularly in the post-2000 period, reflecting divergent country-level dynamics, including both steady improvements and notable declines. Field level data revealed substantial variability as well, with minimum bunch weights ranging from 1.0–24.4 kg and maximum weights 10.9–110 kg for banana, and minimum bunch weights ranging from 1.2-16.6 kg and maximum weights 11-43 kg for plantain (Fig. 2C-D). This likely reflects differences in cultivar types, planting densities, environmental conditions, soil fertility, management practices, and inter-experiment variation across studies.

**Fig. 2.**
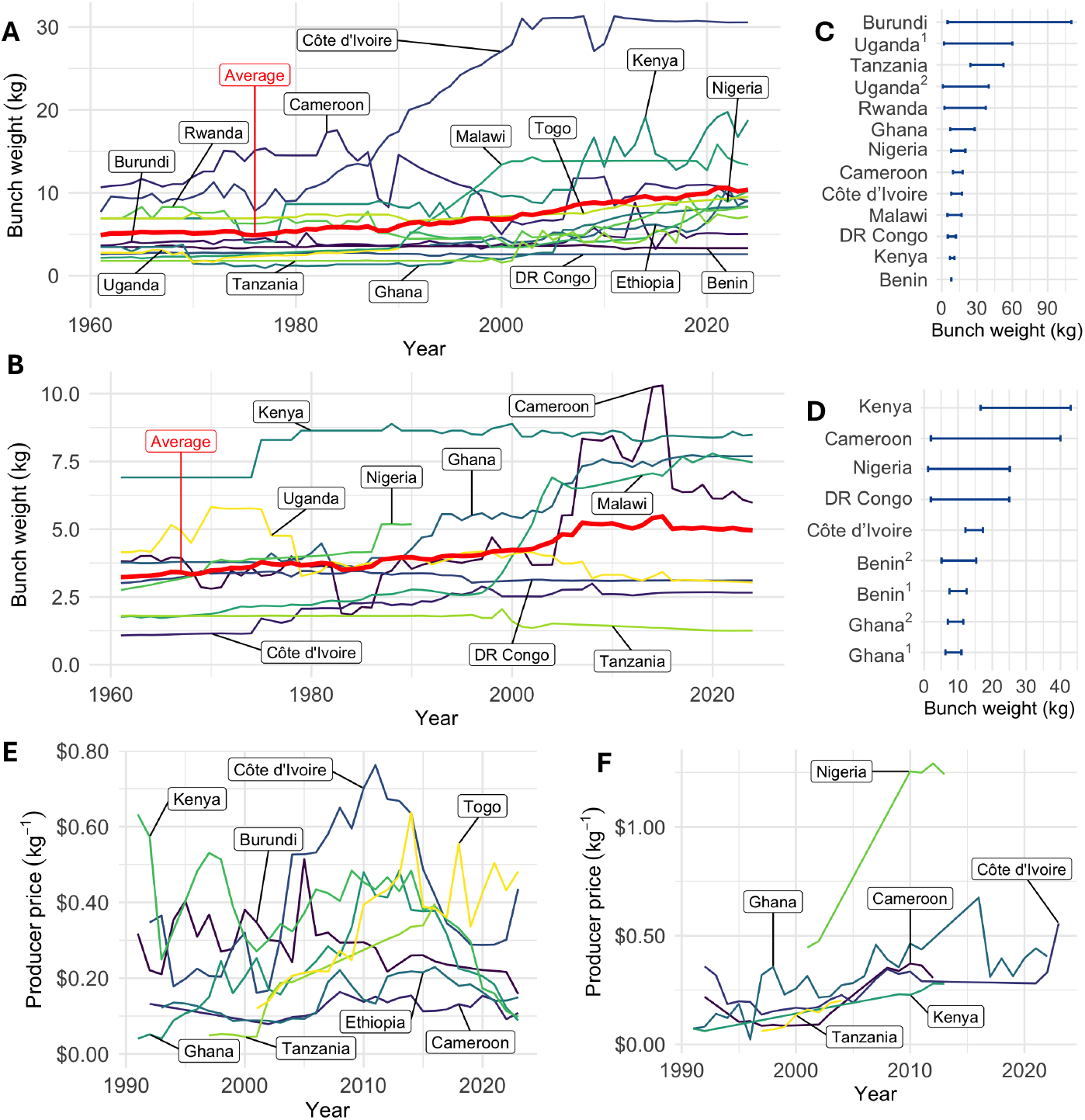
Economic data related to banana and plantain production. (A) FAOSTAT data on banana yield from countries in sub-Saharan Africa [34]. (B) FAOSTAT data on plantain and cooking banana yield from countries in sub-Saharan Africa [34]. (C) Weight of banana bunch from field studies in Benin [35], Burundi [36], Cameroon [37], Côte d’Ivoire [38], Democratic Republic of Congo [39], Ghana [40], Kenya [41], Malawi [42], Nigeria [43], Rwanda [44], Tanzania [45], and Uganda [46, 47]. (D) Weight of plantain bunch from field studies in Benin [48, 49], Cameroon [50], Côte d’Ivoire [51], DR Congo [52], Ghana [53, 54], Nigeria [55], and Kenya [56]. (E) Producer prices (US$) for banana from available countries in sub-Saharan Africa [57]. (F) Producer prices (US$) for plantain from available countries in Sub-Saharan Africa [57].Monthlyuarterly

Clear clusters emerged across countries based on long-term price behaviour (Fig. 2D-F). Burundi, Cameroon, and Ethiopia maintained persistently low and relatively stable producer prices (0.08–0.23 US$/kg). Ghana and Kenya displayed initially high and volatile prices, followed by a pronounced decline and convergence toward 0.09–0.18 US$/kg by 2023. Côte d’Ivoire and Togo showed substantially higher and more variable price regimes, with peaks of 0.76 and 0.64 US$/kg in the mid-2000s before stabilizing in the 2020s. Tanzania exhibited a distinct monotonic increase, rising from 0.045–0.05 US$/kg in the late 1990s to approximately 0.40 US$/kg by 2016, indicating a long-term strengthening of producer value. Plantain price patterns were broadly similar but more volatile and, in some cases, reached higher absolute levels. Cameroon remained in a relatively low-to-moderate price range but showed a clear upward shift after 2008 (*∼*0.30–0.37 US$/kg). Ghana exhibited strong fluctuations, ranging from very low values in the 1990s to peaks above 0.45 US$/kg and a maximum of 0.68 US$/kg in 2016, before stabilizing at intermediate levels. Côte d’Ivoire followed a steady increasing trajectory, reaching 0.56 US$/kg by 2023. Kenya showed gradual strengthening from 0.06–0.07 US$/kg in the early 1990s to 0.25 US$/kg after 2010, while Tanzania also exhibited a consistent upward trend, albeit at lower levels. In contrast, Nigeria formed a distinct high-price cluster, with plantain prices exceeding 1.2 US$/kg in the early 2010s. Overall, while both crops show convergence toward higher producer prices over time, plantain markets are characterized by greater variability and more pronounced country-level divergence.

These national and field-level data were used to define realistic parameter ranges for bunch weight (1–20 kg for banana; 1–17 kg for plantain) and producer prices (0.1–0.8 US$/kg) used in the economic models of BBTV management.

### 2.3 Economic consequences of disease management strategies

Reliable and comparable data on the hourly cost of labour in sub-Saharan Africa are scarce. To reflect realistic economic conditions, we assumed labour costs ranging from 0.5 to 3.0 US$ per hour, consistent with country-specific minimum hourly wages in 2025 (e.g., 0.53 US$ (75 KSH) in Kenya ; 0.27 US$ (718 TZS) in Tanzania; 0.38 US$ (7.15 ZMW) in Zambia, [58]). For analysis purposes, inspection time was set at one minute per plant (*T*_*s*_ = 1*/*60 h), and removal time at 15 minutes per infected plant (*T*_*r*_ = 1*/*4 h). We also consider a zero labour cost scenario to reflect smallholder systems where family labour is not monetised, and inspection and roguing are integrated into routine crop husbandry.

We explored how farm-level economics respond to variations in bunch weight per healthy plant (1–20 kg) and banana price (0.1–0.8 US$); these ranges were informed by the analysis in subsection 2.2. BBTV infection results in no bunch production; therefore, we set bunch weight to zero for exposed and infected plants. Farmer net revenue (FNR) was highly sensitive to inspection frequency and BBTD detection probability (Fig. 3). The figures illustrate three key economic thresholds: (i) parameter combinations where FNR is negative, i.e. banana production is not sustainable, (ii) FNR falls below 1,000 US$, representing conditions in which surveillance is economically marginal, and (iii) factors yielding FNR = 10, 000 US$, reflecting robust profitability. Across both thresholds, economic viability is strongly influenced by labour cost, inspection frequency, and detection probability. At the low-return threshold (1,000 US$), only low labour costs (no labour cost or 0.5 US$/h) allow surveillance to approach marginal profitability, with high-frequency inspections and perfect detection slightly improving revenue.

**Fig. 3.**
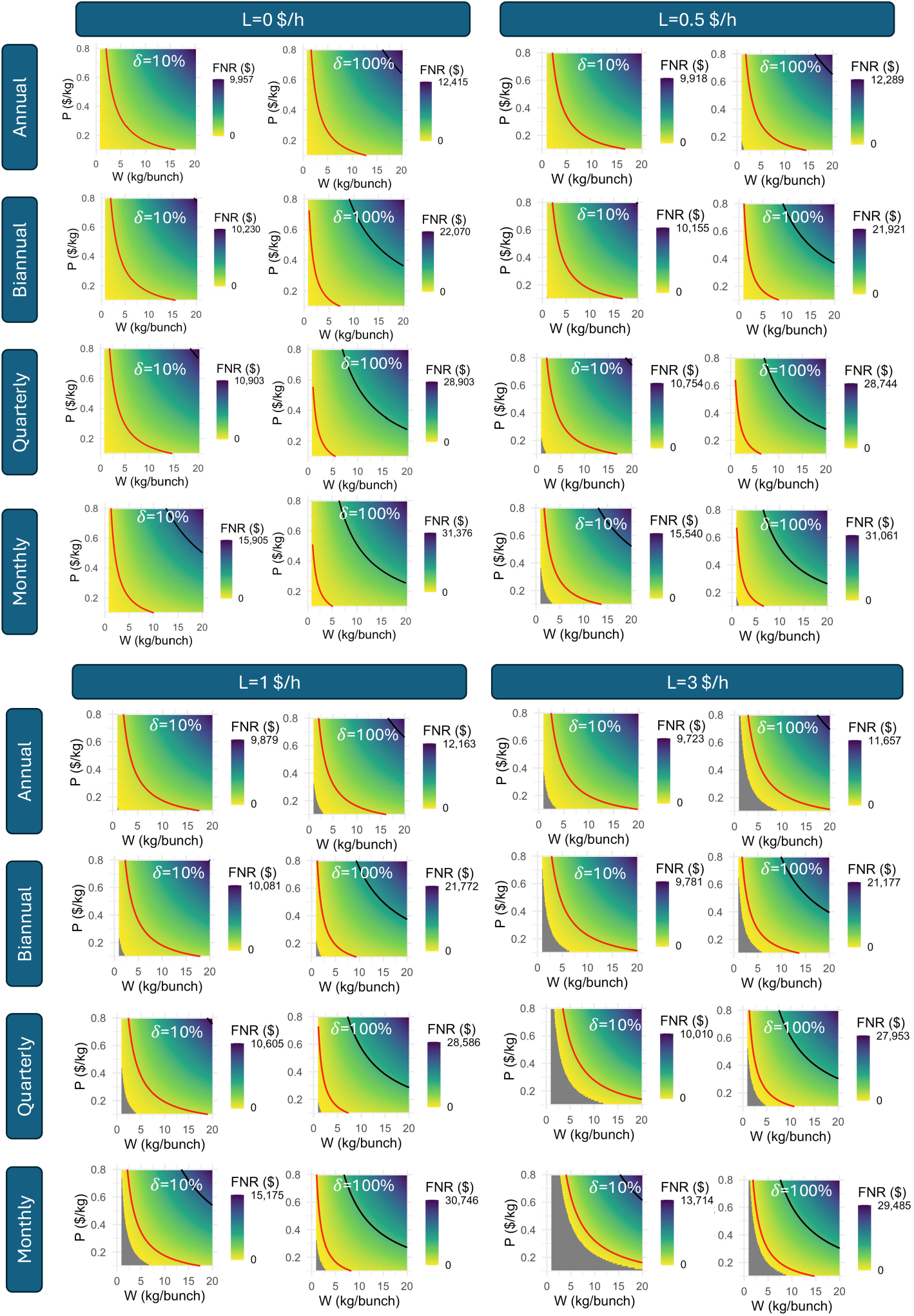
Four-year cumulative farmer net revenue (FNR) as a function of labour cost (*L*, columns), inspection frequency (rows), bunch weight (*W*), banana price (*P*), and BBTD detection probability (*δ*). Dark gray shading (lower left where present) shows area where FNR *≤*0 US$, red curve identifies contour where FNR = 1, 000 US$, and black curve identifies contour where FNR = 10, 000 US$. Calculations are based on an assumption that a farmer requires 1 minute to inspect a plant and 15 minutes to remove an infected plant.

At the higher-return threshold (10,000 US$), surveillance becomes economically justified across a wider range of inspection frequencies and detection probabilities, particularly when labour costs are moderate, and banana prices or bunch weights are high. Low detection probability (10%) with annual inspections requires large bunch weights (up to 57 kg) at low banana prices (0.22–0.23 US$/kg) to reach target profits. As bunch weights decrease, higher banana prices (0.50–0.80 US$/kg) are necessary. Increasing detection probability to 100% reduces reliance on extreme bunch weights or high prices, because effective disease management limits yield losses. More frequent inspections (quarterly or monthly) consistently reduce the banana price required to meet target profits, particularly for intermediate bunch sizes.

Across scenarios where FNR was positive at a detection probability of 10%, increasing the detection probability to 100% substantially enhanced profitability. The mean FNR increase was 110%, with a 95% confidence interval of 74–147%, indicating that effective detection can more than double farm revenue. Importantly, higher detection efficiency not only boosts profits but also stabilizes them, mitigating the impact of price fluctuations or smaller-than-expected bunch weights due to variability in rainfall, temperature, or other abiotic stressors.

### 2.4 Economic optimisation of inspection frequency

To move beyond scenario-specific cost–benefit comparisons and provide operational guidance for smallholder decision-making, we conducted a formal economic optimisation of inspection frequency across epidemiological and market gradients. While previous sections quantified how inspection frequency and detection probability affect BBTV prevalence and farm-level net revenue, farmers and policymakers require clear decision rules specifying the biological and economic conditions under which annual, biannual, quarterly, or monthly inspection is justified, or when no inspection is economically rational. For each parameter combination, we identified the inspection frequency that maximised expected net revenue, constraining outcomes to zero when surveillance was not economically viable.

Optimal inspection frequency and profitability depended on detection probability, bunch weight from healthy plants, banana price, and labour cost (Fig.4). At low price (*P* = 0.25 US$/kg) and low labour cost (*L* = 0.5 US$/h), annual inspection was optimal for small bunches (*≤*4 Kg) at low detection probabilities, with profits up to 83 US$. At higher detection probabilities (*≥*0.7) or larger bunch weights, quarterly or monthly inspection became optimal, and profits increased sharply, reaching 9,204 US$ at 20 kg and *δ* = 1. Increasing labour cost (*L* = 1–3 US$/h) reduced profitability for small bunches, whereas larger bunches still justified inspection, with the optimal frequency that maximised FNR shifting from annual at low detection probability to monthly or quarterly at higher detection probability. For example, at 10 kg and *δ* = 1, quarterly inspection yielded 3,938 US$ (*L* = 1) and 7,876 US$ (*L* = 2). At 3 kg and above, monthly inspection dominated across nearly all detection probabilities, with profits reaching several thousand US$ at higher weights and high detection probability.

**Fig. 4.**
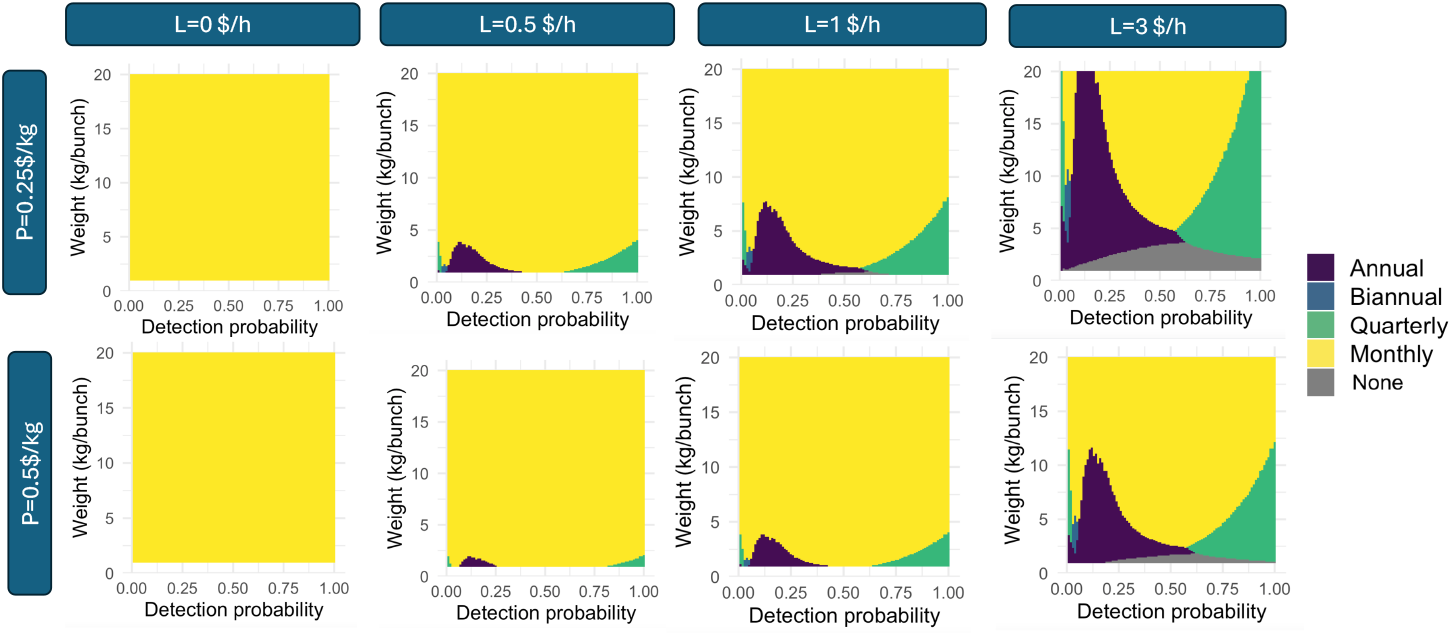
Optimal inspection frequency by detection probability, bunch weight, labour cost (*L*), and price (*P*). Colours show the inspection frequency that maximises expected net revenue over four years: annual (purple), biannual (blue), quarterly (green), monthly (yellow), or none, i.e. FNR *≤*0 (grey). Calculations are based on an assumption that a farmer requires 1 minute to inspect a plant and 15 minutes to remove an infected plant.

Importantly, when labour cost is set to zero - where family labour is not valued as a cash expense — monthly inspection becomes optimal across all bunch weights and detection probabilities. In this case, the trade-off between inspection effort and yield loss is eliminated, and the highest possible surveillance frequency consistently maximises net revenue.

### 2.5 Decision-support tool for optimising farm-level interventions

To support economically informed decision-making for BBTD management, we developed an interactive web application that integrates pre-computed epidemiological simulations with an economic optimisation framework. The tool allows farmers and stakeholders to explore trade-offs between inspection frequency, detection efficiency, and economic outcomes under user-defined production and cost scenarios. The interface, implemented using the **Shiny** package in R (R Core Team), enables real-time recalculation of outcomes across management scenarios. A screenshot of the application is shown in Figure 5. The tool is available in both English and French, making it accessible to a wide range of users in West Africa, where multilingual support is essential for practical adoption and informed decision-making at the farm and community levels.

**Fig. 5.**
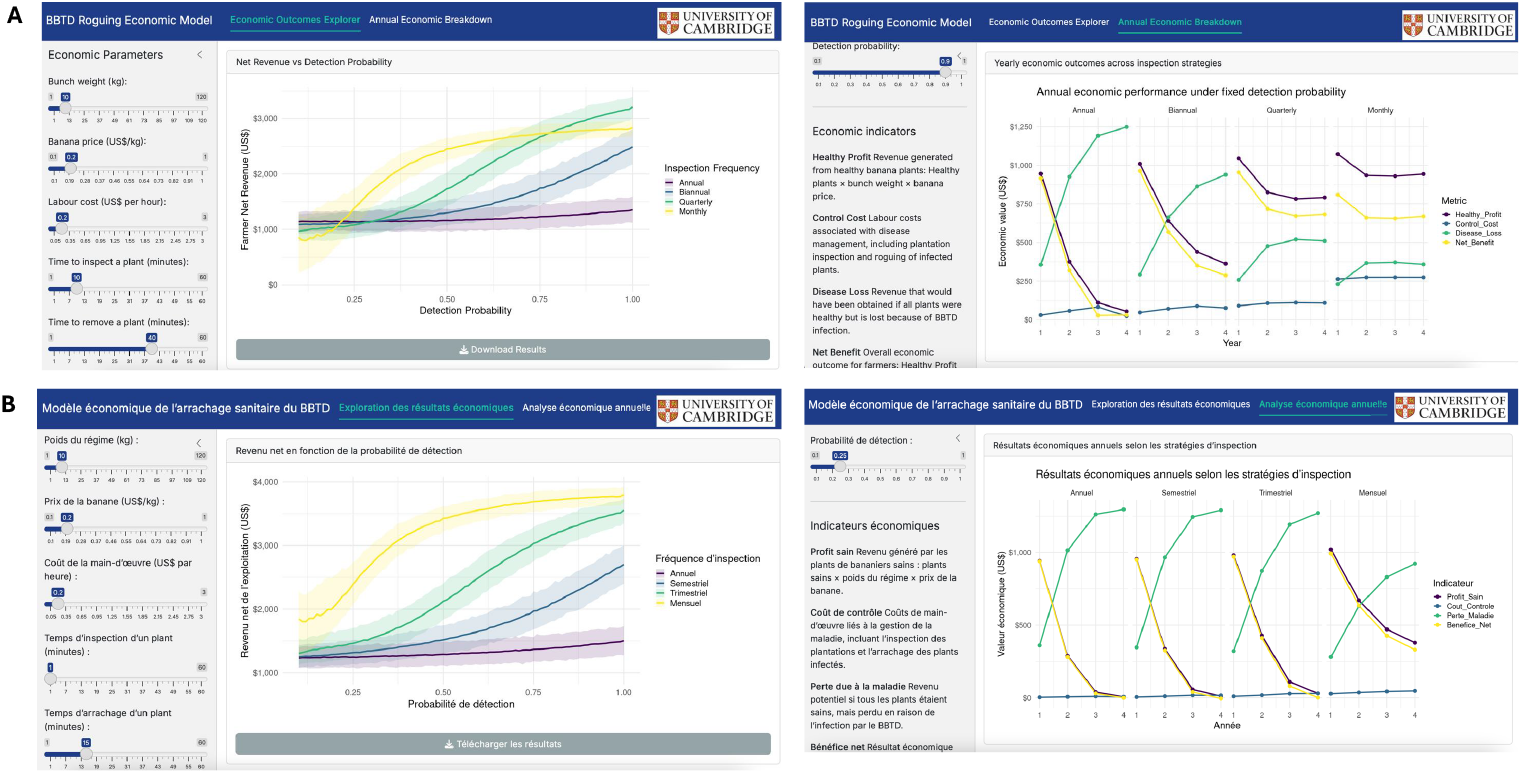
Interactive decision-support tool interface to compute four year Farmer Net Revenue and annual economic outcomes: (A) English version, (B) French version.

The tool evaluates four alternative inspection frequencies: annual, biannual, quarterly, and monthly. For each inspection frequency, detection probability is varied continuously from 0.10 to 1.00 in increments of 0.01. For every combination of inspection frequency and detection probability, the model computes (i) mean net revenue (summed over four years) and (ii) the 95% uncertainty interval (2.5th–97.5th percentile). Outputs are visualised as continuous response curves showing expected profitability as a function of detection probability for each inspection regime, enabling direct comparison of economic performance across surveillance strategies. The interface further allows users to adjust key economic and labour inputs via slider controls. User-defined parameters include bunch weight (1–120 kg), banana price (0.1–1.0 US$/kg), labour cost (0.05–3.0 US$/h), time required to inspect a plant (1–60 minutes), and time required to remove an infected plant (1–60 minutes).

The application facilitates identification of inspection strategies that maximise expected net revenue over the initial period of four years while explicitly accounting for uncertainty in epidemiological outcomes. Because the disease simulations are pre-computed, the platform performs rapid economic recalculations without requiring on-the-fly epidemiological modelling, thereby ensuring responsiveness and usability during stakeholder consultations. By linking epidemiological model outputs with formal economic metrics, the tool provides a transparent and analytically consistent framework for evaluating cost-effectiveness and guiding disease management recommendations at both farm and extension levels. Importantly, the interactive structure enables stakeholders to explore key trade-offs inherent in BBTD management strategies in real time, improving understanding of how surveillance frequency, detection performance, labour costs, and yield parameters jointly determine profitability.

### 2.6 Socio-behavioural determinants of farmer adoption of roguing

Using the dataset described in [59], we fitted a multivariate logistic regression model to identify factors associated with farmer adoption of roguing. Explanatory variables included farmer socio-demographic characteristics, farm and production attributes, and indicators capturing disease awareness, knowledge, and symptom recognition (Appendix Table A1). The model demonstrated good discriminatory performance, with an area under the ROC curve of 0.81 (Appendix Fig. A1).

Few predictors displayed strong statistical associations with roguing behaviour, suggesting which factors might be targeted to encourage uptake of this management practice. Age emerged as one influential factor: younger farmers were substantially less likely to uproot infected plants compared with those over 60 years of age. Farmers aged 20–40 had the lowest odds (*β* = *−*1.99, *p <* 0.05), while those aged 40–60 showed a similar, though weaker, negative association (*β* =*−*1.76, *p <* 0.1). Plantation size had a marginal effect, with farmers managing plots larger than 1, 000 m^2^ exhibiting slightly higher odds (*β* = 1.22, *p <* 0.1).

Diagnostic ability was the only knowledge-related factor significantly associated with roguing: farmers who could correctly recognise BBTD symptoms in the field were more likely to remove infected plants (*β* = 0.94, *p <* 0.05). No significant effects were identified for gender, marital status, ethnicity, education, production orientation, or seed source. Overall, the results indicate that roguing behaviour is strongly associated with farmers’ symptom-recognition capacity, but may also be influenced by economic priorities and cropping system characteristics that shape inspection frequency and incentives.

## 3 Discussion

Recovering and sustaining banana production in regions where BBTV is endemic presents a fundamental challenge for food security in sub-Saharan Africa [60–62]. Because bananas function simultaneously as a staple food, a cash crop, and a cultural asset, disease-driven yield collapse has implications that extend beyond farm income to household nutrition and to the resilience of regional markets. Here, we have integrated epidemiological dynamics, farm-level economics, and behavioural determinants within a single quantitative framework to explore how a single initial introduction of virus-free planting material, followed by routine surveillance, roguing of symptomatic plants, and replanting with suckers taken from surviving field plants, can transition smallholder banana production to a longer-term sustainable system.

The epidemiological results illustrate how inspection frequency and detection probability interact non-linearly to affect disease prevalence. Annual inspection, roguing and replanting with suckers originating in the field—even with high detection accuracy—was insufficient to prevent substantial reinfection, highlighting the persistence of transmission pressure both within the field and from the surrounding endemic landscape. Quarterly or monthly inspections markedly reduced prevalence, and monthly inspection achieved near-suppression even under moderate detection performance. This was reflected in the availability of BBTV-free suckers. For monthly inspections with a moderate detection probability, the proportion of BBTV-free suckers remained above 75% over four years; in contrast, even perfect detection under annual inspections could keep the fraction of BBTV-free suckers only at around 30%. These results demonstrate that frequent inspections act as a strong epidemiological control lever, buffering the system against reinfection-driven degradation of planting material, directly sustaining the informal seed system and reducing the need for repeated external clean-seed inputs.

A simple economic analysis shows that this non-linear interaction between inspection frequency and detection probability translates directly into livelihood outcomes. Profitability is highly sensitive to labour cost, bunch weight, banana price, inspection frequency, and detection probability. Improving detection efficiency from low (10%) to high (100%) more than doubled mean farmer net revenue under many realistic scenarios. Frequent inspection—often perceived as prohibitively labour-intensive—becomes economically rational when detection accuracy is sufficient to prevent cumulative yield loss. Conversely, poor detection erodes the economic case for surveillance even in favourable markets. Both epidemiologically informed control and farmer ability to identify the disease are drivers of profitability.

Importantly, our results illustrate that improving detection probability—for example, through farmer training in symptom recognition—enhances both epidemiological and economic outcomes across nearly all scenarios. When detection probability is low, even frequent inspections may fail to justify their cost under high labour costs and low banana prices, and attempting to eliminate residual infection through continued roguing can reduce net revenue. Under such parameter regimes, maintaining very low endemic prevalence (e.g., 5–10%) may maximise farmer efficiency by balancing inspection costs against avoided yield loss. However, when detection probability is sufficiently high—achievable through targeted training and extension support—the economically optimal outcome shifts toward near-eradication, as the marginal cost of removing the last infected plants is outweighed by the value of saved yield. Here, for simplicity, and in the absence of detailed data, we have assumed that efficiency for BBTV detection is constant throughout the life of the plant. In practice, detection efficiency may vary with plant age. The policy objective should therefore be framed not merely as optimising inspection frequency and replacement planting under resource constraints, but also as investing in farmer diagnostic capacity to raise detection probability, thereby making more intensive management strategies both effective and profitable. Higher detection efficiency consistently expands the range of conditions under which sustained surveillance and roguing outperform less intensive approaches, reinforcing the central importance of farmer training in symptom recognition.

Under climate change, projected banana yields across many African countries are expected to rise by 3–7 t/ha by 2050 [63], while the risk of BBTD is predicted to increase across Central-East Africa (including Uganda and Tanzania), North-East Africa (including Ethiopia), and countries along the 5–10°N latitude band (Côte d’Ivoire, Ghana, Togo, Benin, and Nigeria) [8]. Realizing these potential productivity gains will therefore depend on effective management of BBTV and other pathogens, making disease control essential to safeguarding yield improvements and preventing avoidable losses.

Beyond bananas and plantain, this framework yields transferable insights for other vegetatively propagated crops in which periodic clean-seed interventions [64] or integrated seed health strategies are deployed to limit degeneration and pathogen accumulation [65]. By explicitly quantifying trade-offs among management intensity, detection accuracy, and profitability, it provides a basis for designing interventions that are both economically viable and locally appropriate. The results are intended to inform growers, policymakers, extension services, and development agencies aiming to strengthen the resilience of smallholder systems while reducing dependence on repeated external inputs. They also contribute to ongoing debates on the sustainability of clean-seed programmes in informal seed systems [66, 67]. In much of Africa, informal exchange remains the primary pathway for planting material movement—not only in banana [59] but also in crops such as cassava, yam, and sweet potato [68–72] - highlighting the broader relevance of these findings. Without consistent on-farm monitoring, gains from clean-seed introduction are likely to be short-lived due to reinfection. The analysis demonstrates that combining clean-seed use with systematic roguing and managed sucker propagation can stabilise production, provided detection probabilities exceed economically viable thresholds. Long-term impact therefore hinges not only on external seed supply but also on strengthening local surveillance and improving the performance of informal seed networks, where skilled farmers or regulated multipliers can act as reliable sources of low-risk planting material.

The socio-behavioural analysis from Benin reinforces this dimension. The strongest predictor of roguing adoption was farmers’ ability to recognise BBTV symptoms, while broader socio-economic characteristics were largely insignificant. Management behaviour is therefore constrained primarily by diagnostic competence rather than structural demographics. In the Benin data, younger farmers were less likely to rogue infected plants, suggesting that intergenerational knowledge transfer and targeted training may be critical for continuity. Similar findings were reported in Uganda, where participatory awareness campaigns led to high levels of symptom recognition and knowledge of disease transmission; farmers who correctly identified banana bacterial wilt symptoms were significantly more likely to adopt recommended control measures [73]. These results position symptom recognition as a pivotal node linking epidemiology, economics, and behaviour within the production system.

From a food-systems perspective, epidemiological shocks propagate through production losses to influence caloric availability and price stability. In countries where banana provides a substantial share of daily caloric intake, sustained yield loss would reduce food availability and amplify local market volatility. Because bananas are largely domestically consumed rather than export-oriented in much of sub-Saharan Africa, production shocks transmit rapidly to household welfare. By quantifying economic and epidemiological parameters jointly, we illustrate how operational guidance for designing interventions can enhance both productivity and resilience. Investments that improve detection capacity—through farmer training, extension support, or digital diagnostics—are necessary complements to clean-seed distribution and to sustaining the effectiveness of subsidised seed programmes.

In conclusion, our study advances an integrated quantitative framework linking epidemiology, economics, and behaviour in smallholder banana systems. Clean-seed programmes can succeed in endemic BBTV settings—but only when paired with sufficiently frequent inspection and high detection accuracy. More broadly, the framework demonstrates how disease management in vegetatively propagated crops should be conceptualised as a dynamic optimisation problem under biological and economic constraints. Sustainable banana production depends on strategies to ensure clean or low risk planting material, which can be achieved by equipping farmers with the skills and incentives to detect and remove infection early. Strengthening local surveillance capacity thus emerges as a cornerstone of household daily nutrition and food security.

## 4 Materials and methods

### 4.1 Protocol for re-establishing smallholder banana production

To develop best-practice guidance for introducing clean banana planting material into BBTV-affected areas of sub-Saharan Africa, we constructed a modelling framework to evaluate how surveillance and roguing can limit disease establishment and persistence within a representative single-field system. We assumed that farmers begin with *N* units of clean planting material that are 100% virus-free. Banana plants are arranged on a regular grid (3 m between and 2.5 m within rows) within this single representative field configured as 30 rows by 20 columns, comprising a total of 651 plants. This is equivalent to a density of 1447 plants *ha*^*−*1^, and corresponds to a plot size of 0.45 ha, consistent with typical banana field sizes in Africa, which range from 0.01 ha to 1 ha, with an average of 0.2 ha [74]. We consider a monocropped field planted with a single variety, located in an endemic area and continually exposed to a high risk of infection from the surrounding landscape in which there are infected banana plants in the surrounding areas, for example in previously infected abandoned banana crops. We assume that the monocropped filed has previously been cleared of banana plants before replanting.

### 4.2 BBTV transmission and management simulation

We simulate BBTV spread using a stochastic, individual-based model that captures both spatial and temporal dynamics. Each banana plant is assigned to one of five mutually exclusive states: susceptible (S), exposed (E), infectious (I), symptomatic and detected (D), or removed (R). A banana plant in the exposed state has already been infected with Banana bunchy top virus—typically through feeding by a viruliferous aphid—but is not yet capable of transmitting the virus to other plants [75]. Plant transitions to the infectious state once viral titers in the phloem are high enough for aphids to acquire and subsequently transmit the virus [76]. The framework accommodates both immediate removals upon detection and delayed removals that may occur due to logistical constraints after inspection. Removed plants are replaced with suckers sourced from the “healthy-looking” remaining plants in the field, which the farmer assumes to be healthy, and it is assumed that a sufficient number of such plants are available at any time during the period of the model run.

BBTV transmission into a field occurs via two distinct pathways: primary infection, in which the virus is introduced from external sources either by migrating viruliferous aphids or the introduction of infected planting materials, and secondary infection, representing transmission between infectious and susceptible plants within the field by aphids through feeding. The infection pressure experienced by a susceptible plant *i* at time *t* is described by:

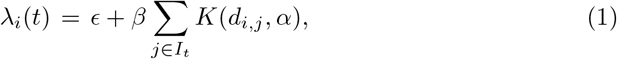

where *ϵ* is the primary infection rate, *β* is the secondary infection rate, *K*(*d, α*) is the dispersal kernel for secondary infection, *d*_*i,j*_ is the distance between plant *i* and plant *j*, and *I*_*t*_, is an index set of infectious plants at time *t*. We define the dispersal kernel as:

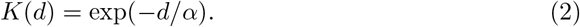

The latent period of BBTD corresponds to the delay between infection by banana bunchy top virus and the onset of detectable symptoms or infectiousness. This delay is not fixed and can vary with environmental conditions and plant growth dynamics. In particular, it is often related to the rate of leaf production, as symptom expression is associated with newly emerged leaves [77]. Consequently, the latent period may be represented as a function of leaf emergence rate (LER), which itself exhibits seasonal variation and can be expressed as a function of day of the year, following [78]:

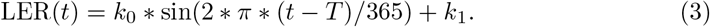

Here *k*_0_ represents the amplitude of seasonal variation in leaf emergence, *k*_1_ is the mean (baseline) leaf emergence rate, and *T* is a phase-shift parameter that determines the timing of the seasonal peak in LER within the annual cycle. In other words, *T* controls when during the year leaf emergence reaches its maximum, effectively aligning the periodic function with local climatic seasonality. We used values for the parameters, *k*_0_ = 0.056, *k*_1_ = 0.062, from [78], while setting *T* = 90 days to align peak leaf emergence with the onset of the rainy season in West Africa, which is a key driver of banana growth dynamics [79]. The latent period of BBTV was parameterised following the experimental analysis of Allen [31], who estimated it as the time required for the production of approximately 3.7 banana leaves after infection. This estimate was derived from observed relationships between infection timing, leaf emergence, and the subsequent appearance of symptoms, and reflects the coupling between host growth dynamics and symptom development driven by systemic accumulation of BBTV.

We assumed that *p*% of suckers produced by exposed plants are infectious, and that 100% of suckers produced by infectious plants are infectious. Suckers are not tested for BBTV before planting, therefore representing an additional source of infection. We further assumed that all infected plants eventually express symptoms and will ultimately be detected and removed. The model has been parameterised in [80], including estimates for infection rates, aphid dispersal parameters, and proportion of infected suckers re-planted in the field.

We simulated inspection, detection and roguing as follows. Symptomatic plants were rogued according to an inspection cycle for all plants in each field. The quality of inspection was directly dependent on the probability of detecting a symptomatic plant. During model simulations, all infected plants were ordered according to the day they became infectious, and then a proportion of plants equal to the detection probability was removed according to this order. We investigated values of detection probabilities in the range from 10% to 100% with steps of 10%. Lower values of detection probability correspond to surveys conducted by less experienced observers (e.g., farmers), where symptom recognition is more uncertain and only a subset of symptomatic plants may be identified, particularly given variability in symptom expression and cases requiring expert diagnosis. Higher probabilities represent surveys conducted by trained inspectors with greater ability to detect and correctly identify symptomatic plants. We varied the frequency of inspection, with inspection taking place at a regular interval of one month (monthly), three months (quarterly), six months (biannually) or twelve months (annually). We ran each scenario for 1000 simulations. When simulating detection of less than 100%, we sampled plants which were infectious with a probability equivalent to the difference between the time of inspection and the time they became infectious.

### 4.3 Cost–benefit analysis of BBTV management

We developed a plant-level cost–benefit framework to evaluate the economic performance of surveillance and roguing as management strategies for BBTV. We defined the farmer net revenue (FNR) in year *y* as the revenue from healthy plants minus the costs of surveillance and roguing:

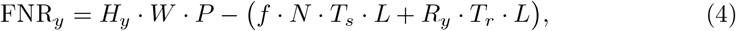

where *N* is number of plants in the field, *H*_*y*_ is the number of healthy plants which were planted at least one year ago [81], *R*_*y*_ is number of removed plants, *W* is the bunch weight, *P* is the banana price, *f* is number of surveys per year, *T*_*s*_ and *T*_*r*_ are the labour time per inspection and per removal, respectively, and *L* is the hourly cost of farm labour. Total farmer net revenue was obtained by summing the annual *FNR*_*y*_ over all years. The number of total healthy and removed plants were obtained by taking average over the simulations at the end of each year.

### 4.4 Economic input data

National-level yield data for desert banana, and plantain/cooking banana (1961–2024) were obtained from the FAOSTAT database of the Food and Agriculture Organization of the United Nations [34]. To contextualize these large-scale statistics within the production realities of smallholder farmers in sub-Saharan Africa, we expressed national yield estimates (in tonnes per hectare, *tha*^*−*1^) as an equivalent weight per individual banana bunch, using the field-scale configuration described in Section 2.1.

To complement national banana production statistics, we compiled field-level data from the peer-reviewed literature. This effort identified 13 studies reporting banana production metrics across 12 countries in Sub-Saharan Africa: Benin [35], Burundi [36], Cameroon [37], Côte d’Ivoire [38], Democratic Republic of Congo [39], Ghana [40], Kenya [41], Malawi [42], Nigeria [43], Rwanda [44], Tanzania [45], and Uganda [46, 47]. For plantain, we retrieved nine studies providing bunch weight data from Benin [48, 49], Cameroon [50], Côte d’Ivoire [51], DR Congo [52], Ghana [53, 54], Nigeria [55], and Kenya [56].

Producer price data (1990–2025) were sourced from the HDX platform [57], which collates FAOSTAT-derived market information curated by the World Food Programme. FAO reports producer prices that represent what farmers receive at the farm gate (the point where the crop leaves the farm).

### 4.5 Survey on BBTD management practices

The data were obtained from [59], which reports information collected from trained farmers in Benin between October 2018 and June 2021. A questionnaire was used to record information on farmer characteristics, production practices, knowledge about disease, and the implementation of various management methods. Factors influencing farmers’ decisions to adopt specific management practices were inferred using logistic regression.

## Author contributions

R.R., A.B.O., J.E.T., C.S., and C.A.G. designed the study; R.R., J.E.T. and C.A.G. initiated and planned the manuscript; R.R. developed the epidemiological model, performed simulations and analysis, and implemented the Shiny application; R.R., A.B.O., M.Z.-T., U.R.A., C.S., P.L.K., J.E.T. and C.A.G. wrote the manuscript.

## Data availability

The datasets and code for this study can be found in the Gtihub repository https://github.com/rretkute/BBTVModelSmallFieldData.

## Appendix A Logistic regression for roguing uptake

**Table A1.**
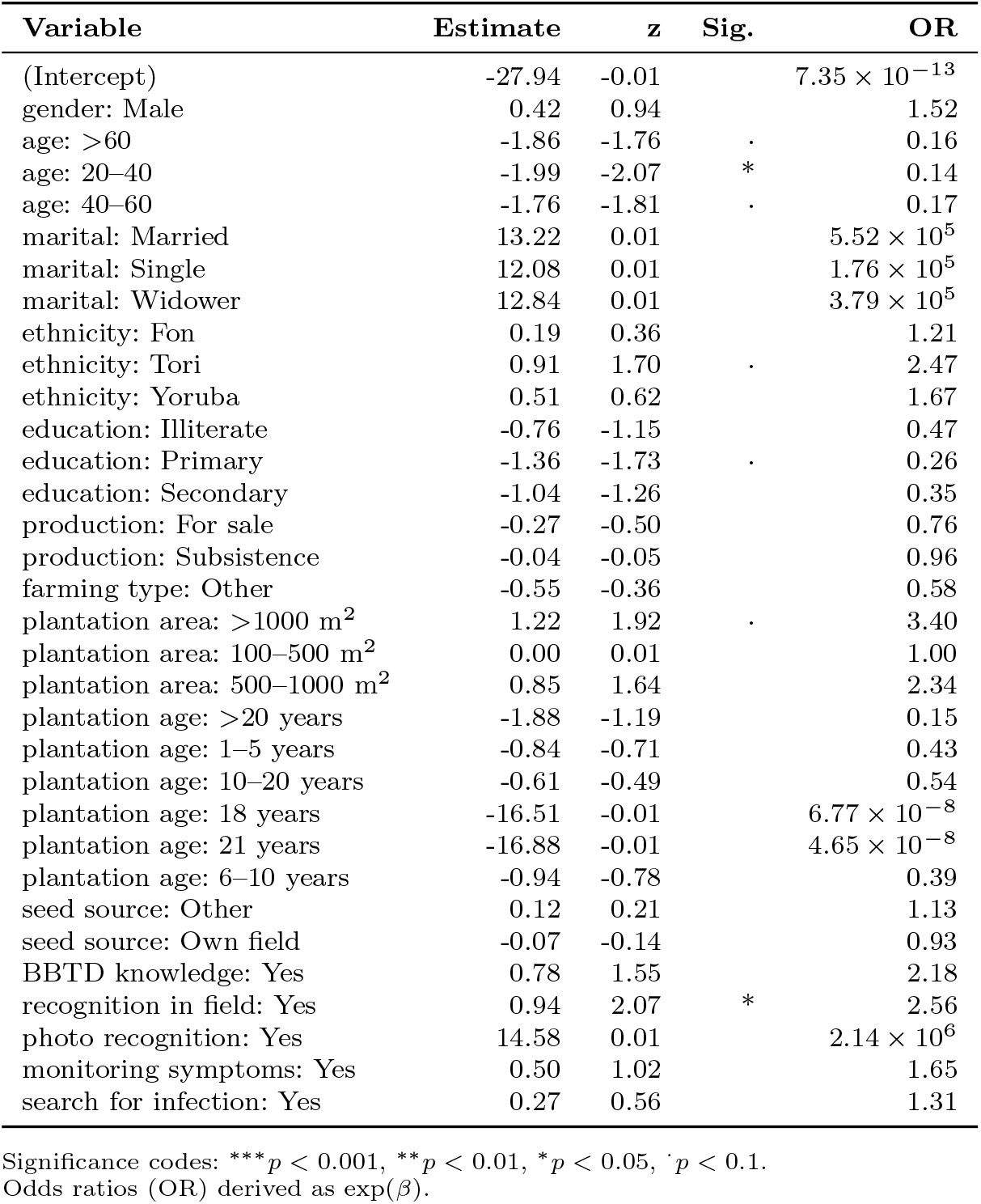
Determinants of farmer implementation of roguing: logistic regression coefficients and odds ratios.

| Variable | Estimate | z | Sig. | OR |
| --- | --- | --- | --- | --- |
| (Intercept) | -27.94 | -0.01 | | $7.35 \times 10^{-13}$ |
| gender: Male | 0.42 | 0.94 |  | 1.52 |
| age: >60 | -1.86 | -1.76 | . | 0.16 |
| age: 20–40 | -1.99 | -2.07 | * | 0.14 |
| age: 40–60 | -1.76 | -1.81 | . | 0.17 |
| marital: Married | 13.22 | 0.01 | | $5.52 \times 10^5$ |
| marital: Single | 12.08 | 0.01 | | $1.76 \times 10^5$ |
| marital: Widower | 12.84 | 0.01 | | $3.79 \times 10^5$ |
| ethnicity: Fon | 0.19 | 0.36 |  | 1.21 |
| ethnicity: Tori | 0.91 | 1.70 | . | 2.47 |
| ethnicity: Yoruba | 0.51 | 0.62 |  | 1.67 |
| education: Illiterate | -0.76 | -1.15 |  | 0.47 |
| education: Primary | -1.36 | -1.73 | . | 0.26 |
| education: Secondary | -1.04 | -1.26 |  | 0.35 |
| production: For sale | -0.27 | -0.50 |  | 0.76 |
| production: Subsistence | -0.04 | -0.05 |  | 0.96 |
| farming type: Other | -0.55 | -0.36 |  | 0.58 |
| plantation area: >1000 m<sup>2</sup> | 1.22 | 1.92 | . | 3.40 |
| plantation area: 100–500 m<sup>2</sup> | 0.00 | 0.01 |  | 1.00 |
| plantation area: 500–1000 m<sup>2</sup> | 0.85 | 1.64 |  | 2.34 |
| plantation age: >20 years | -1.88 | -1.19 |  | 0.15 |
| plantation age: 1–5 years | -0.84 | -0.71 |  | 0.43 |
| plantation age: 10–20 years | -0.61 | -0.49 |  | 0.54 |
| plantation age: 18 years | -16.51 | -0.01 | | $6.77 \times 10^{-8}$ |
| plantation age: 21 years | -16.88 | -0.01 | | $4.65 \times 10^{-8}$ |
| plantation age: 6–10 years | -0.94 | -0.78 |  | 0.39 |
| seed source: Other | 0.12 | 0.21 |  | 1.13 |
| seed source: Own field | -0.07 | -0.14 |  | 0.93 |
| BBTD knowledge: Yes | 0.78 | 1.55 |  | 2.18 |
| recognition in field: Yes | 0.94 | 2.07 | * | 2.56 |
| photo recognition: Yes | 14.58 | 0.01 | | $2.14 \times 10^6$ |
| monitoring symptoms: Yes | 0.50 | 1.02 |  | 1.65 |
| search for infection: Yes | 0.27 | 0.56 |  | 1.31 |

Significance codes: \*\*\* $p < 0.001$ , \*\* $p < 0.01$ , \* $p < 0.05$ , . $p < 0.1$ . Odds ratios (OR) derived as $\exp(\beta)$ .

---

**Fig. A1.**
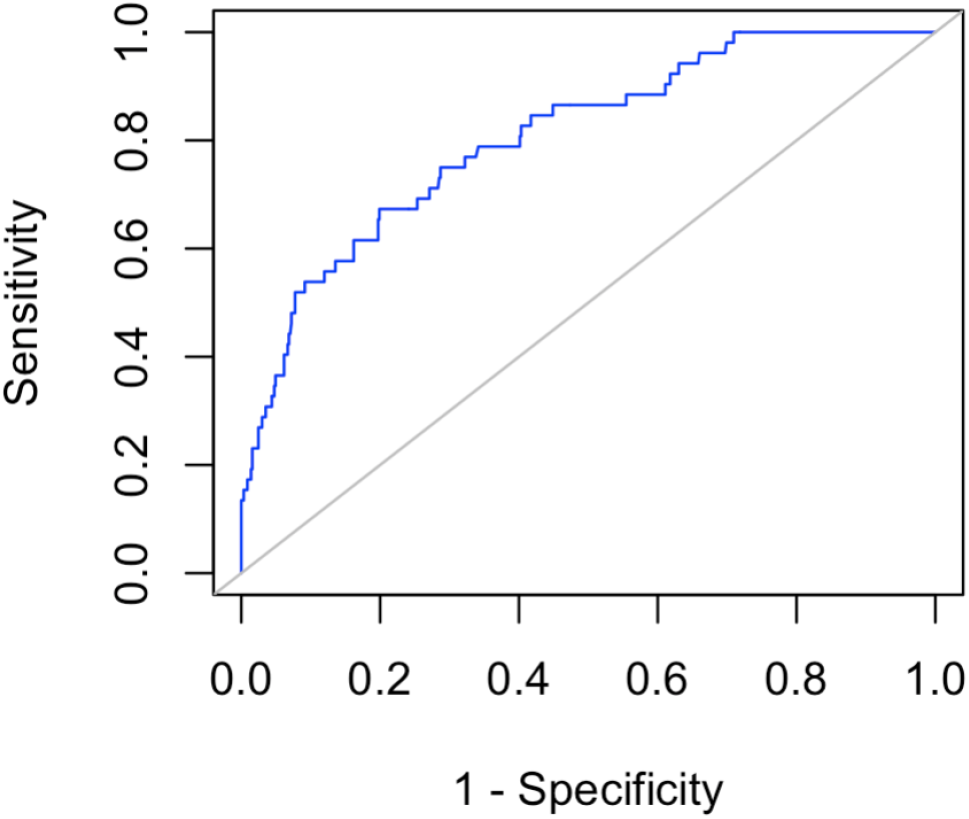
Receiver operating characteristic (ROC) curve for logistic regression model predicting farmer intention to implement roguing as a management of BBTD.

